# Intermediate molecular phenotypes to identify genetic markers of anthracycline-induced cardiotoxicity risk

**DOI:** 10.1101/2023.01.05.522844

**Authors:** Aurora Gómez-Vecino, Roberto Corchado-Cobos, Adrián Blanco-Gómez, Natalia García-Sancha, Sonia Castillo-Lluva, Ana Martín-García, Marina Mendiburu-Eliçabe, Carlos Prieto, Sara Ruiz-Pinto, Guillermo Pita, Alejandro Velasco-Ruiz, Carmen Patino-Alonso, Purificación Galindo-Villardón, María Linarejos Vera-Pedrosa, José Jalife, Jian-Hua Mao, Guillermo Macías de Plasencia, Andrés Castellanos-Martín, María del Mar Sáez Freire, Susana Fraile-Martín, Telmo Rodrigues-Teixeira, Carmen García-Macías, Julie Milena Galvis-Jiménez, Asunción García-Sánchez, María Isidoro-García, Manuel Fuentes, María Begoña García-Cenador, Francisco Javier García-Criado, Juan Luis García, María Ángeles Hernández-García, Juan Jesús Cruz Hernández, César Augusto Rodríguez-Sánchez, Alejandro Martín-Ruiz, Estefanía Pérez-López, Antonio Pérez-Martínez, Federico Gutiérrez-Larraya, Antonio J. Cartón, José Ángel García-Sáenz, Ana Patiño-García, Miguel Martín, Teresa Alonso Gordoa, Christof Vulsteke, Lieselot Croes, Sigrid Hatse, Thomas Van Brussel, Diether Lambrechts, Hans Wildiers, Chang Hang, Marina Holgado-Madruga, Anna González-Neira, Pedro L Sánchez, Jesús Pérez Losada

## Abstract

Cardiotoxicity due to anthracyclines (CDA) affects cancer patients, but we cannot predict who may suffer from this complication. CDA is a complex disease whose polygenic component is mainly unidentified. We propose that levels of intermediate molecular phenotypes in the myocardium associated with histopathological damage could explain CDA susceptibility; so that variants of genes encoding these intermediate molecular phenotypes could identify patients susceptible to this complication. A genetically heterogeneous cohort of mice generated by backcrossing (N = 165) was treated with doxorubicin and docetaxel. Cardiac histopathological damage was measured by fibrosis and cardiomyocyte size by an Ariol slide scanner. We determine intramyocardial levels of intermediate molecular phenotypes of CDA associated with histopathological damage and quantitative trait loci (ipQTLs) linked to them. These ipQTLs seem to contribute to the missing heritability of CDA because they improve the heritability explained by QTL directly linked to CDA (cda-QTLs) through genetic models. Genes encoding these molecular subphenotypes were evaluated as genetic markers of CDA in three cancer patient cohorts (N = 517) whose cardiac damage was quantified by echocardiography or Cardiac Magnetic Resonance. Many SNPs associated with CDA were found using genetic models. LASSO multivariate regression identified two risk score models, one for pediatric cancer patients and the other for women with breast cancer. Molecular intermediate phenotypes associated with heart damage can identify genetic markers of CDA risk, thereby allowing a more personalized patient management. A similar strategy could be applied to identify genetic markers of other complex trait diseases.

## Introduction

Cardiotoxicity due to anthracyclines (CDA) is a frequent problem in cancer patients that limits the efficacy of chemotherapy (Caron and Nohria 2018). Long-term cardiotoxicity has repercussions for oncological disease prognosis (Patnaik et al. 2011) and a far-reaching impact on patients’ quality of life (Pein et al. 2004). Anthracyclines produce acute necrosis and apoptosis of cardiomyocytes, leading to myocardial fibrosis and varying degrees of chronic functional damage, and even heart failure (Chatterjee et al. 2010). The degree of chronic CDA depends on many factors, including dose, age, gender, previous heart diseases, and combined treatment with other drugs (Grenier and Lipshultz 1998). Since it is difficult to determine which patients will develop chronic CDA, efforts have been made to identify genetic risk markers. However, current evidence about the role of pharmacogenomic screening in anthracycline therapy is mixed because of the heterogeneity of the results obtained so far (Leong et al. 2017).

The diversity of the results observed when attempting to identify CDA genetic markers may result from the CDA being a complex polygenic disease or complex trait influenced by multiple genes contributing at systemic, tissue, cellular, and molecular levels (Duan et al. 2007). However, it is difficult to determine how best to measure the genetic influence of complex traits. The proportion of phenotypic variation explained by the genetic component is known as narrow-sense heritability. Though, the genetic variants associated with complex diseases account for only 10-20% of the phenotypic variation attributable to genetics. The genetic variants responsible for the remaining phenotypic variation cannot be identified and are considered missing heritability; identifying its origins remains a contentious matter (Manolio et al. 2009).

Complex diseases and traits arise from multiple intermediate phenotypes or endophenotypes that participate in their pathogenesis (Blanco-Gomez et al. 2016). Also,–intermediate phenotypes may themselves be complex traits. For instance, myocardial infarction is a complex-trait disease, susceptibility to which is determined by intermediate phenotypes such as arterial hypertension, hypercholesterolemia, and the response to tobacco exposure. However, these intermediate phenotypes are also complex traits influenced by lower-ranking intermediate phenotypes located at the systemic, cellular, and molecular levels. Genetic determinants directly act on the intermediate molecular phenotypes implicit in this multidirectional network of intermediate phenotypes (Blanco-Gomez et al. 2016). Indeed, a specific protein and RNA would be the simplest intermediate phenotypes, controlled by a few genes (including the coding gene, the genes encoding promoter-regulating transcription factors, and those controlling post-translational activity regulators, such as phosphorylation) (Civelek and Lusis 2014). The variable phenotypic presentation of complex diseases is related to the expressivity of their intermediate phenotypes (Gottesman and Gould 2003; Blanco-Gomez et al. 2016). Thus, different degrees of susceptibility to CDA could result from the variable expression of intermediate molecular phenotypes participating in CDA pathogenesis.

It was previously proposed that the missing heritability of complex traits could be due to genes that exert their influence at the level of intermediate phenotypes, such as arterial hypertension. However, they would not be powerful enough to be detected at the level of the primary complex phenotype, such as acute myocardial infarction (Castellanos-Martin et al. 2015; Blanco-Gomez et al. 2016), in whose pathogenesis they participate. This possibility is consistent with the heritability being missing because differences between many common variants cannot attain statistical significance in GWAS studies (International Schizophrenia et al. 2009; Yang et al. 2010; Loh et al. 2015; Shi et al. 2016). Therefore, it was hypothesized that genes lacking sufficient strength to be detected at the main trait level (as mediated pleiotropy of intermediate phenotypes) account for a proportion of the missing heritability of this complex trait (Blanco-Gomez et al. 2016).

It is predicted that common small-effect genes affecting a complex trait might be located throughout much of the genome. For example, between 71% and 100% of 1 Mb windows could contribute to the heritability of schizophrenia (Loh et al. 2015), and thousands of eQTLs may control blood gene expression (Vosa et al. 2021). Additionally, mathematical models predict that between 0.1 and 1% of SNPs have a causal effect on most diseases studied (Khera et al. 2018). These observations align with the omnigenic model that purports to explain missing heritability (Boyle et al. 2017; Liu et al. 2019). Thus, attributing missing heritability to the genetic determinants that influence numerous intermediate phenotypes of a complex disease would require thousands of genes acting on its phenotypic variation and susceptibility (Blanco-Gomez et al. 2016). However, it is difficult to identify some of these genetic markers that also would help predict the risk of complex trait diseases. Efforts have been made to develop genome-wide polygenic scores based on thousands of genetic variants (Khera et al. 2018).

In this work, we propose that genetic determinants linked to intermediate molecular phenotypes of CDA could help quantify susceptibility to this chemotherapy complication. We illustrate the rationale of our study in **Fig. 1**.

**Figure 1.**
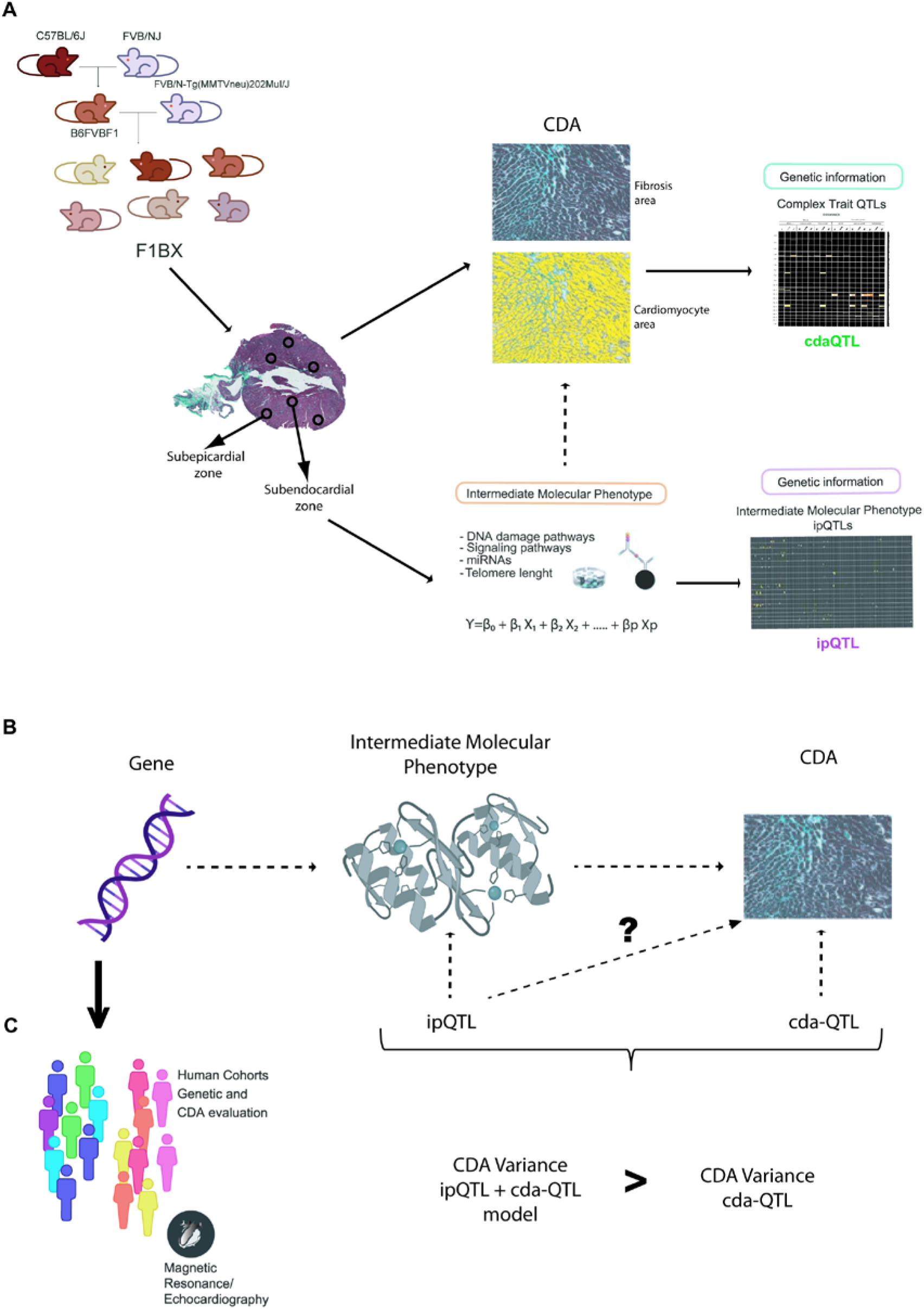
Scheme of the general approach. **A**) This paper studies the different susceptibility to anthracycline cardiotoxicity (CDA) in a cohort of mice generated by backcrossing. The degree of CDA was quantified at the histopathological level using an Ariol slide scanner in the subepicardial and subendocardial zones of the heart. In addition, the levels in the myocardium of different molecules (intermediate phenotypes) associated with cardiotoxicity were quantified. Subsequently, genetic regions or quantitative trait loci (QTL) associated with CDA (cdaQTL) and other QTLs associated with intramyocardial levels of intermediate molecular phenotypes (ipQTL) were identified. **B**) We then evaluate whether some of the ipQTLs contribute significantly to the phenotypic variation (susceptibility) of CDA, despite not being directly linked to it. To do this, we evaluate using genetic models whether the phenotypic variation explained by the ipQTL together with the cdaQTL exceeded that explained by the cdaQTL alone. If so, the genetic determinant linked to the myocardial levels of this intermediate molecular phenotype contributes to CDA susceptibility. **C**) The genes responsible for ipQTLs are unknown; however, in principle, any genetic determinant that influences the levels of these molecules associated with cardiotoxicity could contribute to CDA susceptibility. Therefore, allelic forms of the encoding genes of these intermediate molecular phenotypes would be candidates to be evaluated in the human population.

Therefore, we assess the degree of CDA in a cohort of genetically heterogeneous mice generated by backcrossing. To identify intermediate molecular phenotypes of CDA, we quantify the myocardial levels of signaling pathways, microRNAs (miRNAs), and telomere length and evaluate their association with histopathological heart damage after chemotherapy. We then demonstrate that the genetic determinants associated with the levels of some of these intermediate molecular phenotypes in the myocardium contribute to the heritability of CDA. To do so, (i) first, we identify quantitative trait loci (QTLs) linked to the CDA intermediate molecular phenotypes (ipQTLs); (ii) second, we show that ipQTLs integrated into genetic models with a QTL directly linked to myocardial damage (cda-QTL) explain a more significant proportion of the phenotypic variability than does the cda-QTL alone. We conclude that since ipQTLs are not directly linked to CDA, they must contribute to its missing heritability. Thus, genetic determinants influencing the levels of intermediate molecular phenotypes in the myocardium, including the genes encoding the intermediate molecular phenotypes themselves, would contribute to the heritability of CDA. Subsequently, genes encoding intermediate molecular phenotypes of CDA may be markers of susceptibility to this side effect of chemotherapy. We evaluate this possibility in three cohorts of human patients.

## Results

### Cardiotoxicity due to anthracyclines behaves as a complex trait in a genetically heterogeneous mouse cohort

CDA is a complex trait, and as such, identifying the genetic component in humans that influences CDA susceptibility is a challenging task (Leong et al. 2017). However, crosses between syngeneic mouse strains enable part of the genetic background components, and thereby complex traits, to be identified (Hunter and Crawford 2008; Castellanos-Martin et al. 2015). Therefore, we generated a cohort of mice with a heterogeneous genetic background by backcrossing to identify genetic determinants linked to CDA susceptibility. We crossed *MMTV-Erbb2/Neu* transgenic mice with FVB background with F1 non-transgenic mice to generate the backcrossed cohort (hereafter, F1BX). These mice were treated with doxorubicin or combined therapy once they had developed breast cancer. Since doxorubicin and docetaxel are used in human cancer chemotherapy (De Laurentiis et al. 2008), the mice were treated once they had developed breast cancer induced by the MMTV-*Erbb2/Neu* transgene (Guy et al. 1992) (**Fig. 2A**). Previous studies showed that, after anthracycline chemotherapy, a subclinical injury could be detected at the histopathological level in the myocardium even before functional damage had occurred (Billingham et al. 1978; Friedman et al. 1978). Anthracyclines induce the death of cardiomyocytes that are replaced by fibrosis, leading to atrophy of the left ventricle. Also, the atrophy of cardiomyocytes secondary to the toxicity of anthracyclines is described, which can be observed early by CMR (Ferreira de Souza et al. 2018). However, in the long term, there is hypertrophy of the remaining cardiomyocytes due to ventricular remodeling secondary to diastolic overload when heart failure occurs (Goorin et al. 1990; Lipshultz et al. 1991; Piek et al. 2016). Thus variable grades of cardiomyocyte hypertrophy, myocytolysis, and fibrosis are characteristic features of ventricular remodeling associated with anthracycline exposure (Segura et al. 2015). Thus, interstitial fibrosis and cardiomyocyte area modification are phenotypes of pathological cardiac remodeling and chronic CDA (Lipshultz et al. 1991; Piek et al. 2016). We quantified both pathophenotypes in the myocardium after chemotherapy using an Ariol slide-scanner to evaluate the degree of CDA, considering the global, subendocardial, and subepicardial zones of the heart (**Fig. 2A** and **Supplemental_Fig_S1.pdf.**)

**Figure 2.**
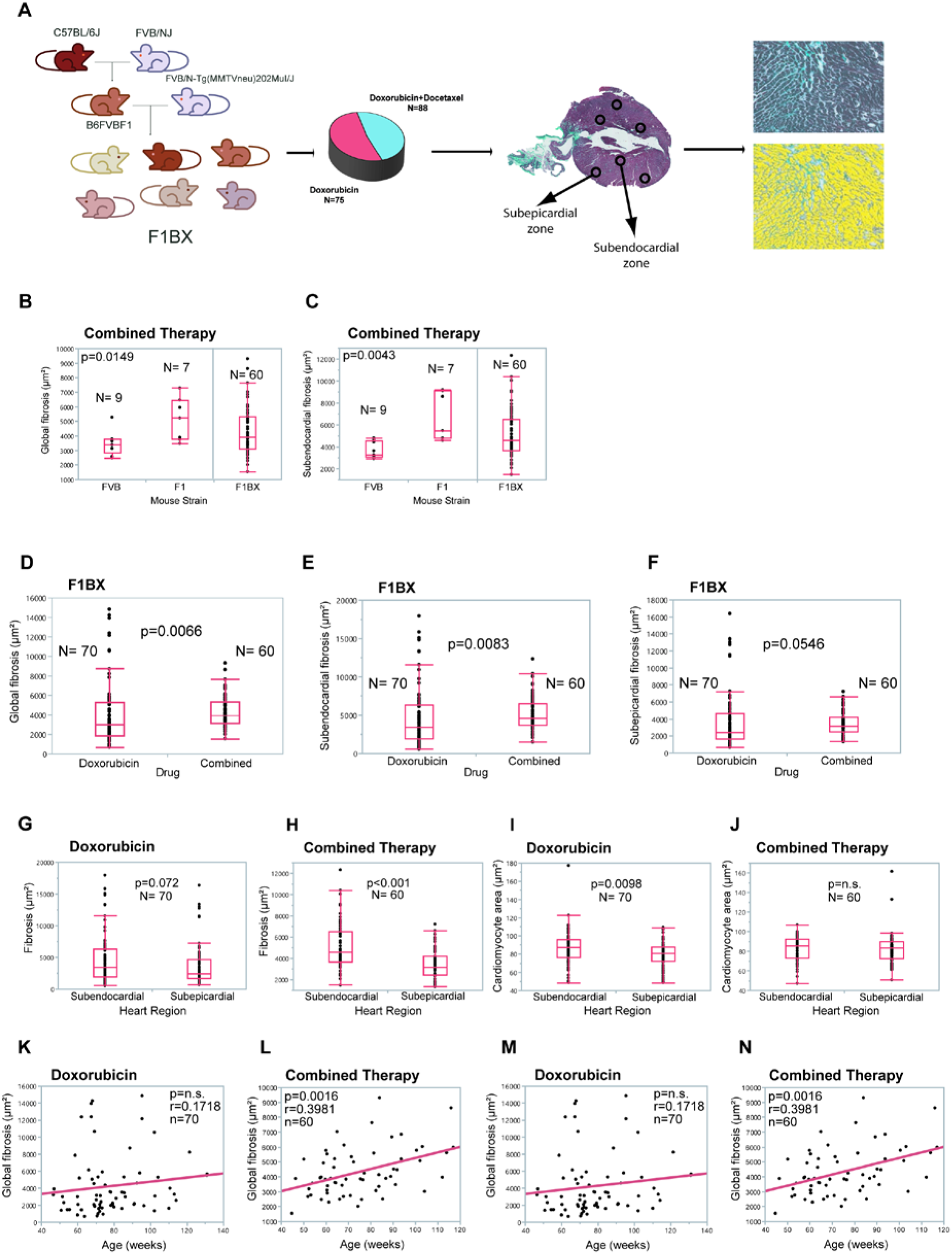
Anthracycline-induced cardiotoxicity differs according to genetic background, therapy regime, and age. **A**) Cardiotoxicity due to anthracyclines (CDA) is evaluated in a cohort of mice generated by backcrossing. The mice are treated with chemotherapy, and cardiotoxicity is quantified at the histopathological level. **B**, **C**) Comparison of cardiotoxicity between mouse strains of homogeneous genetic background. After combined chemotherapy, FVB mice showed less global heart fibrosis (B) and less subendocardial fibrosis (C) than F1 mice. No differences exist between the parental strains after chemotherapy with doxorubicin alone (not shown). Note the distribution of heart fibrosis as a continuum in F1BX mice (A, B). **D-F**) In the genetically heterogeneous cohort of F1BX mice, the combined treatment is more cardiotoxic than doxorubicin alone in terms of the degree of global (D), subendocardial (E), and subepicardial (F) fibrosis. **G-J**) CDA was higher in the subendocardial zone than in the heart’s subepicardial location in F1BX1 mice, in fibrosis (G, H) and cardiomyocyte area (I, J). Mann–Whitney U test. **K**, **L**) CDA in the whole cohort of F1BX mice based on age and type of chemotherapy. Global fibrosis is not correlated with age for the treatment with doxorubicin alone (K); however, it is positively correlated with age after the combined treatment (L), as estimated by the Spearman correlation coefficient. **M**, **N**) Global fibrosis increases with age in old mice treated with doxorubicin (L) or combined therapy (M), as determined by the Spearman correlation coefficient. We show only those results that were statistically significant.

Initially, we explored and compared heart fibrosis and the cardiomyocyte area between FVB and F1 mice. F1 mice treated with combined therapy had significantly higher levels of global (*p* = 0.0149) and subendocardial (*p* = 0.0043) fibrosis than did FVB mice (**Fig. 2B, C**); we did not find more differences between both strains (**Supplemental_Table_S1A.xls.**)

We evaluated CDA in the F1BX genetically heterogeneous cohort of mice generated by backcrossing. As expected, the observed degree of cardiotoxicity spanned a wider range than seen in the parental strains and was distributed as a continuum throughout the F1BX mice (**Fig. 2B, C**), as is characteristic of complex traits(Mackay 2009). We then compared the CDA after doxorubicin treatment and combined therapy in the F1BX mice. Cardiotoxicity was higher after the combined therapy for global (*p* = 0.0066), subendocardial (*p* = 0.0083), and subepicardial fibrosis (*p* = 0.0546) than when doxorubicin was administered alone (**Fig. 2D-F**); we did not observe differences in the cardiomyocyte area between both regimes of therapy (**Supplemental_Table_S1B.xls**). Globally, cardiotoxicity was more significant in the subendocardial than in the subepicardial area of the heart (**Fig. 2G-J**). As expected, the combined therapy was more cardiotoxic than therapy with doxorubicin alone (**Supplemental_Table_S1C.xls**.)

Chronic CDA susceptibility increases with age in humans (Aapro et al. 2011), so we evaluated how heart damage varied with mouse age. Heart fibrosis increased with age after combined therapy (*p* = 0.0016) but not significantly after therapy solely with doxorubicin in global heart and subepicardial and subendocardial zones (**Fig. 2K-N** and **Supplemental_Table_S2A.xls**). We divided the cohort into young and old groups of mice based on the sample’s median age of 71 weeks. Fibrosis increased with age in the old group after doxorubicin alone (*p* = 0.0052) and the combined treatment (*p* = 0.021) but not in young mice (**Supplemental_Table_S2A.xls**). We did not observe differences in cardiomyocyte area (**Supplemental_Table_S2B.xls**). Together, these results indicate that older mice were more sensitive to chronic CDA than younger mice, as previously observed in cancer patients (Aapro et al. 2011).

The behavior and distribution of chronic CDA in F1BX mice as a complex trait are like those found in humans. Indeed, CDA was significantly greater after combined therapy than with doxorubicin alone (Salvatorelli et al. 2006; Salvatorelli et al. 2007) and with increasing age (Aapro et al. 2011; Armenian et al. 2017) and was distributed as a complex trait in the F1BX mice (Mackay 2009). All these similarities justified using the F1BX backcross model to identify the genetic background component linked to chronic CDA.

### Intermediate phenotype levels of molecular origin in the myocardium are associated with chemotherapy-induced cardiotoxicity

CDA is a complex trait. As such, its pathogenesis is influenced by different intermediate phenotypes at the systemic, tissue, cellular and molecular levels (Gianni et al. 2008; Blanco-Gomez et al. 2016). We used this mouse backcross strategy as a simplified model to seek intermediate molecular phenotypes associated with CDA. We quantified the levels of several molecules in the myocardium after chemotherapy and evaluated their association with heart fibrosis and the cardiomyocyte area (**Fig. 3A**). The molecules were selected based on their involvement in the pathogenesis of cardiomyopathy, as described in previous reports. Using multiplex bead arrays, we quantified levels of the myocardium proteins involved in antigenotoxicity pathways (34) and cell-signaling pathways that favor or inhibit heart damage caused by anthracycline (35). We also used qPCR to determine the miRNAs involved in cardiac diseases and cardiotoxicity (Ruggeri et al. 2018) and in controlling myocardium telomere length (De Angelis et al. 2010).

**Figure 3.**
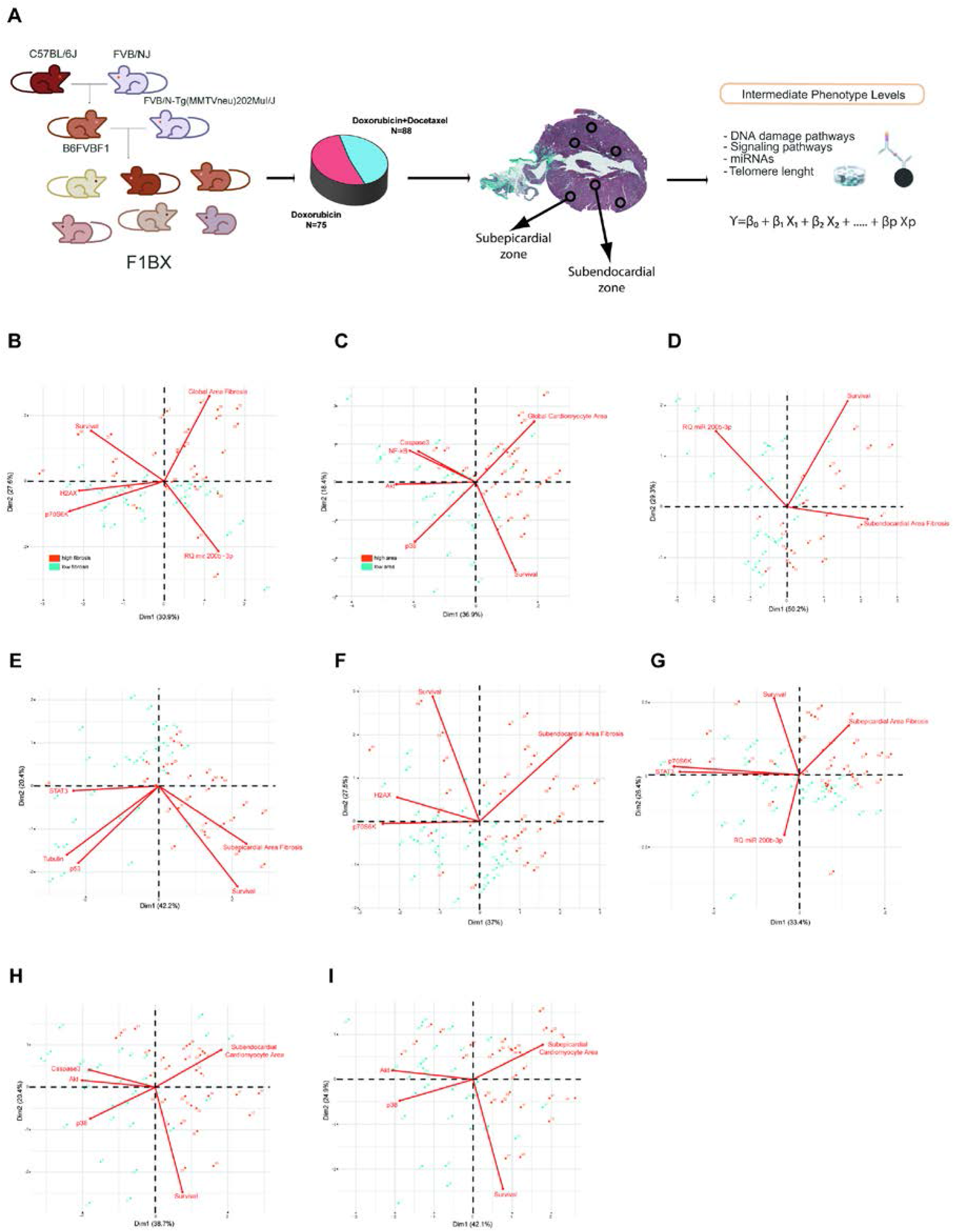
Intermediate molecular phenotypes are associated with cardiotoxicity due to anthracyclines (CDA). **A**) Quantification in the myocardium of F1BX mice of different intermediate molecular phenotypes after chemotherapy. **B-I**) Principal component analyses classify mice with high and low CDA susceptibility based on the levels of intermediate molecular phenotypes in the myocardium under different conditions: fibrosis after doxorubicin (B), cardiomyocyte area after the combined therapy (C), subendocardial zone fibrosis (combined therapy) (D), subepicardial zone fibrosis (combined therapy) (E), subendocardial zone fibrosis (doxorubicin) (F), subepicardial fibrosis zone (doxorubicin) (G), subendocardial cardiomyocyte area after combined therapy (H), subepicardial cardiomyocyte area (combined therapy) (I). Mice with high (brown) and low (blue) levels of fibrosis and cardiomyocyte area were differentiated by the median. This figure is related to Supplemental Table S3.

The levels of intermediate molecular phenotypes involved in the pathogenesis of complex traits should be statistically significantly associated with the complex trait (Gottesman and Gould 2003). Indeed, some intermediate molecular phenotypes were associated with the variation of fibrosis and the cardiomyocyte area in the F1BX mice (**Supplemental_Table_S3.xls**). The integration of these intermediate molecular phenotypes by principal component analyses (PCA) helped to distinguish between mice with high and low CDA in different conditions (**Fig. 3B-I**.)

We subsequently used multiple regression to evaluate which intermediate phenotypes were most important for defining CDA (**Supplemental_Table_S4.xls**). For instance, after doxorubicin chemotherapy, young mice with low levels of P70S6K(pT412) and old mice with low levels of H2AX(pS139) in the myocardium had higher global fibrosis in the heart. After combined therapy, young mice with high levels of CREB1(pS133) presented high global fibrosis in the myocardium. AKT1(pS473), P38MAPK(pT180/pY182), β-tubulin and TP53(S15), and the miRNAs miR210_3p, mR215_5p, Let7d_5d, and Let7d_5p were associated with CDA under a variety of conditions. These selected molecular intermediate phenotypes also help to identify mice with high and low CDA susceptibility by PCA (**Supplemental_Fig_S2.pdf**.)

In summary, it may be concluded that all these molecules associated with heart fibrosis and the cardiomyocyte area after chemotherapy could be intermediate phenotypes related to chronic CDA susceptibility variation in the F1BX cohort of mice.

### Identification of genetic determinants linked to intermediate molecular phenotypes of CDA

It has been indicated that genetic determinants linked to the intermediate phenotypic function of a complex trait could account for some of the phenotypic variations in the latter and contribute to its missing heritability (Gottesman and Gould 2003; Blanco-Gomez et al. 2016). Among the genetic determinants that determine the functional activity of an intermediate molecular phenotype, there are fundamentally those that regulate its levels. These determinants also include the gene that encodes the molecule with its regulatory sequences in *cis* and another series of genes in QTL regions located in *trans* that helps regulate molecular levels and activity (Civelek and Lusis 2014; Vosa et al. 2021), which we call intermediate phenotype QTLs (ipQTLs).

Following on, we asked whether the ipQTLs associated with intermediate molecular phenotypes of CDA contribute to the phenotypic variation of the latter. We set about integrating the ipQTLs with directly linked QTLs into genetic models with CDA (cdaQTLs) to determine whether they could account for more of the CDA phenotype variation than that explained solely by cda-QTLs (**Fig. 1**). Accordingly, we looked for ipQTLs and cdaQTLs in the F1BX genetically heterogeneous mice that could be used subsequently in the genetic models (**Fig. 1**). Thus, firstly, we looked for the genetic regions (ipQTLs) associated with the myocardium levels of the intermediate molecular phenotypes identified (**Fig. 4A**). The global scenario of ipQTLs identified is shown as a heatmap (**Figure 4B**), and the specific information for each genetic locus is presented in **Supplemental_Table_S5.xls**.

**Figure 4.**
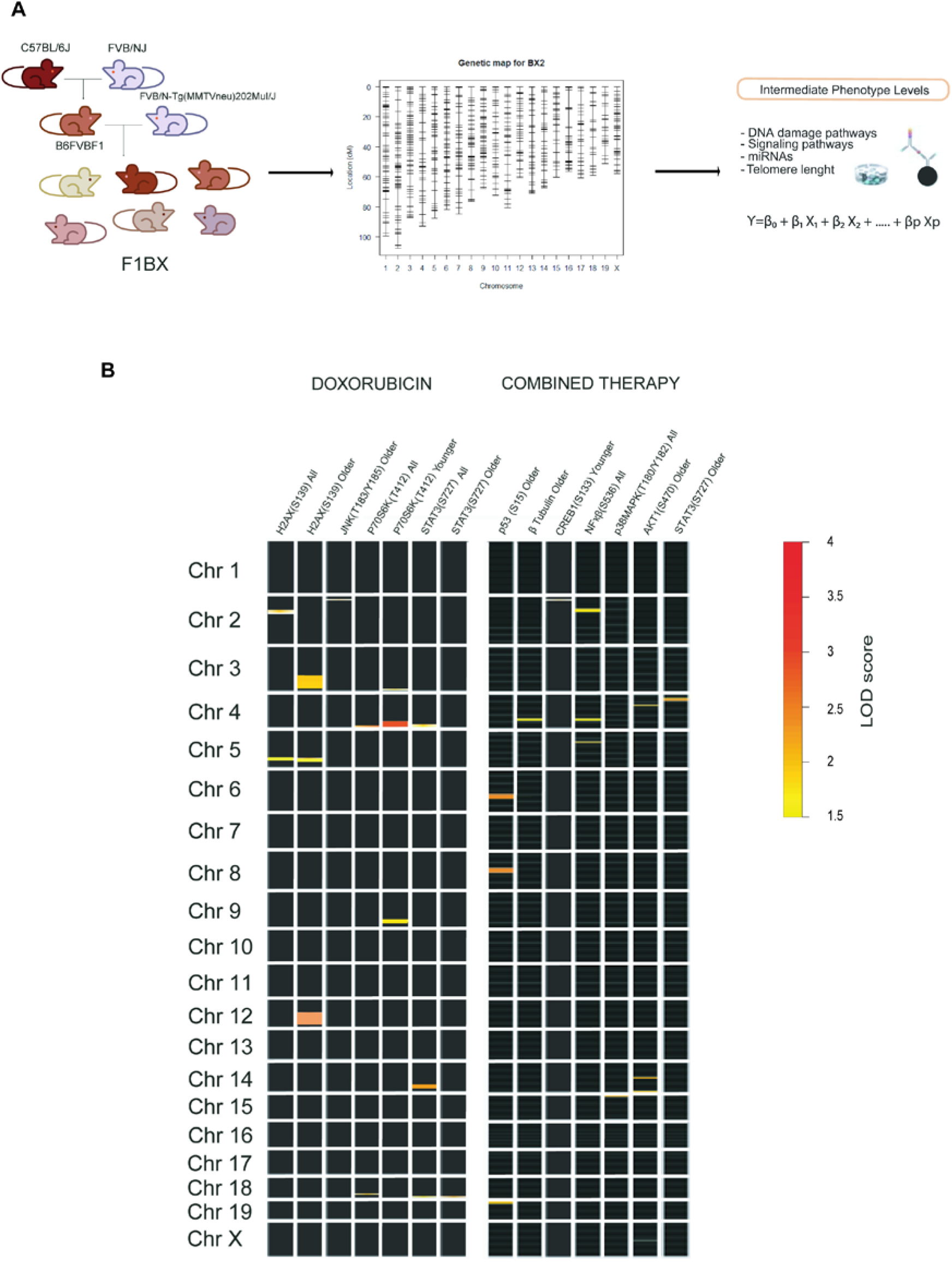
Identification of quantitative trait loci associated with intermediate molecular phenotypes (ipQTLs) of CDA susceptibility. **A**) F1BX1 mice generated by backcrossing were genotyped using 1499 SNPs on an Illumina platform. Linkage analysis was performed to detect ipQTLs. **B**) The heatmap shows the global scenario of ipQTLs after doxorubicin or combined therapy. Each square represents a chromosome; its number is on the left. Each square’s intensity mark signifies the degree of association (LOD score) between the genetic markers and the phenotype according to the indicated scale. The marks’ location of each square occupies the relative area within each chromosome (centromere and telomere positions above and below, respectively). Only linkages with a LOD score > 1.5 (suggestive) are represented. R/qtl software was used to identify ipQTLs. The exact location of each ipQTL in each chromosome and the associated genetic markers are shown in **Supplemental_Table_S5A.xls** (doxorubicin therapy) and **Supplemental_Table_S5B.xls** (combined therapy).

### Identification of genetic determinants directly linked to CDA (cdaQTLs)

Following on, we searched for cdaQTLs associated with heart fibrosis and the cardiomyocyte area based on the type of chemotherapy (anthracycline alone or combined therapy) in different conditions: heart zone (whole heart, subendocardium or subepicardium) and age (young or old mice) (**Fig. 5A**). QTLs linked to CDA under these different conditions were represented as heatmaps to visualize the global scenario of the genetic regions linked to heart damage after doxorubicin or combined therapy (**Fig. 5B, C**, and **Supplemental_Table_S6.xls**.)

**Figure 5.**
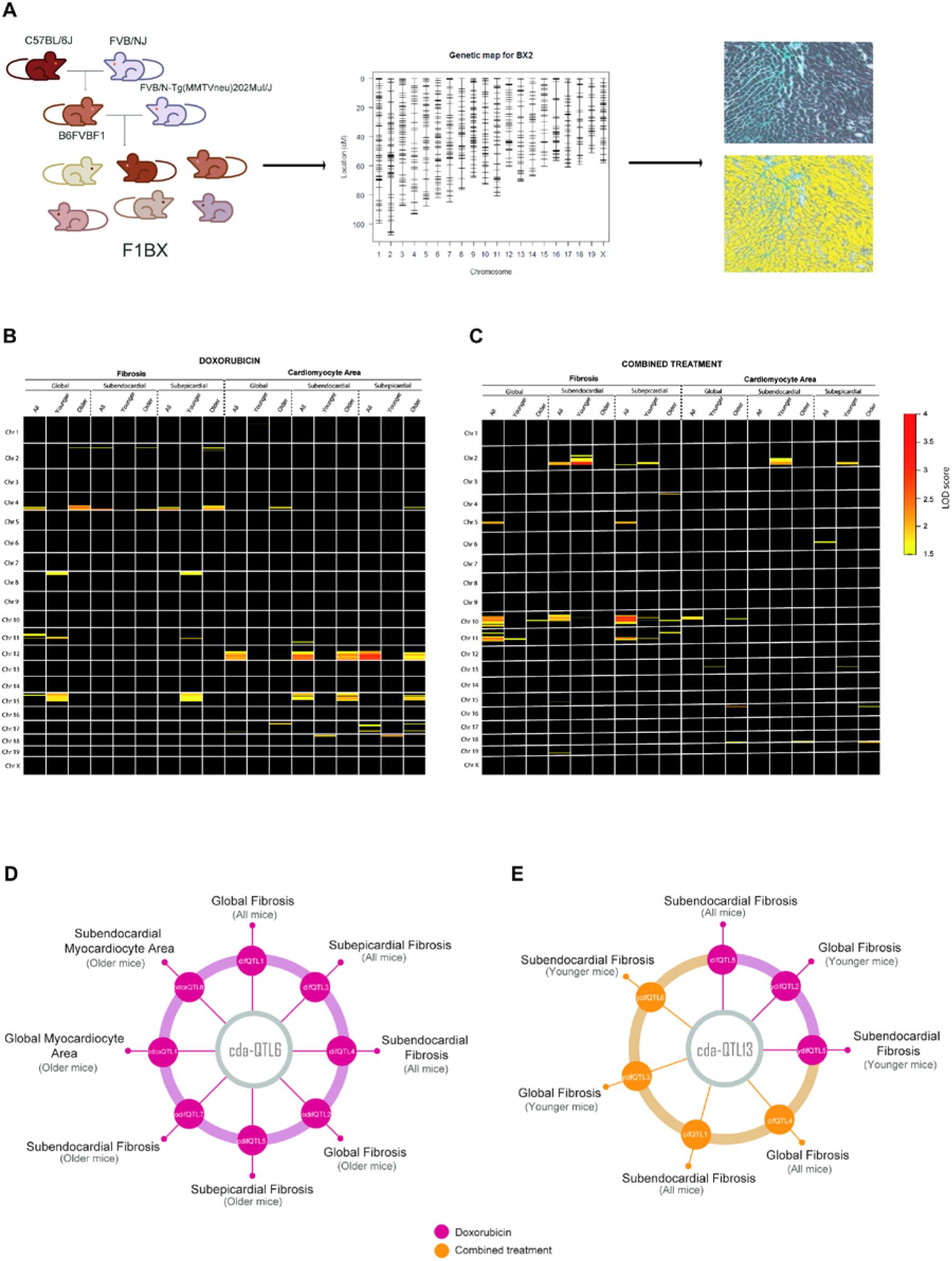
Quantitative trait loci linked to cardiotoxicity due to anthracyclines (cda-QTLs). **A**) The genotyping of the F1BX mouse cohort permitted to locate cdaQTLs directly linked to the CDA susceptibility measured at the histopathological level. **B, C)** The heatmaps show the global scenario of the cda-QTLs linked to cardiac fibrosis and cardiomyocyte area under different conditions (age and type of therapy): after chemotherapy with doxorubicin (B) or after combined treatment (C). For both panels, B and C: only QTLs with a LOD score > 1.5 were plotted along each chromosome. Analyses were carried out with R/qtl software. Each square represents one chromosome; its number is indicated on the left. According to the indicated scale, the intensity of marks within each square indicates the degree of association (LOD score) between the genetic markers and the phenotype. The mark’s location corresponds to its relative location within the chromosome (centromere and telomere positions above and below, respectively). The exact locations of each cda-QTL and its associated genetic markers are shown in Supplemental Table S5. **D**) The cda-QTL6 was associated with CDA under different conditions in old mice. cda-QTL6 colocalized with different QTLs associated with CDA under different conditions in old mice, specifically with difQTL1 (doxorubicin-induced fibrosis QTL1), difQTL3, difQTL4, odifQTL 4 (old mice odifQTL4), odifQTL5, odifQTL7, odcaQTL1 (old mice doxorubicin-induced cardiomyocyte area QTL1) and odcaQTL8. **E**) cda-QTL13 was associated with fibrosis in younger mice and colocalized with the following QTLs: difQTL5, ydifQTL2 (young mice difQTL2), ydifQTL5, cifQTL4 (combined therapy-induced fibrosis QTL4), cifQTL11, ycifQTL3 (young mice QTL3), and ycifQTL6. The numerical data (LOD score and peak locations of markers) from panels C and D are shown in Supplementary Tables S5 and S6.

Eighty cda-QTLs were identified, but some were in the same chromosome (Chr.) and were genetic regions simultaneously linked to several CDA conditions. In the end, we identified 27 cda-QTLs in full (**Supplemental Table S6**). For example, the same QTL was sometimes associated with the degree of fibrosis and the cardiomyocyte area under different conditions; this was the case of cda-QTL6 on Chr. 4 after doxorubicin treatment and cda-QTL11 on Chr. 10 after combined therapy (**Figure 5D** and **Supplemental_Table_S7.xls**). The same QTL and pathophenotype were occasionally associated in both chemotherapy regimens, e.g., the cda-QTL13 on Chr. 11 and heart fibrosis (**Figure 5E** and **Supplemental_Table_S7.xls**). Notably, the cda-QTL6 on Chr. 4 was explicitly associated with CDA in old mice, whereas the cda-QTL13 on Chr. 11 was most frequently related to CDA susceptibility in young mice (**Fig. 5D, E** and **Supplemental_Table_S7.xls**). The identification of multiple QTLs associated with CDA confirmed the polygenic component of susceptibility to this complication, even in a simplified model like that of the F1BX mouse cohort(Duan et al. 2007).

### ipQTLs integrated into genetic models with cdaQTLs account for more phenotypic variation of CDA than explained by cda-QTLs alone

Our next step was to integrate the ipQTLs with cda-QTLs into the genetic models(Broman et al. 2003) to evaluate whether these could account for more of the CDA phenotype variation than that explained solely by the cda-QTL (**Fig. 6A**). In doing so, we wanted to demonstrate that these ipQTLs contribute to the missing heritability of the CDA. The cda-QTL would enable the ipQTLs contributing to the missing heritability of CDA to be revealed by the genetic models (**Fig. 1B**). As examples, we selected cdaQTL6 and cdaQTL13 (**Fig. 5D, E**) to evaluate whether ipQTLs integrated with these cdaQTLs could account for more of the CDA phenotype variation than that explained solely by cdaQTL6 and cdaQTL13 (**Fig. 6A**). First, we estimated the phenotypic variance of global fibrosis attributable to them in the F1BX cohort. CdaQTL6 explained 22.17% of the CDA variance in global fibrosis in old mice after doxorubicin chemotherapy, and cdaQTL13 accounted for 28.82% and 26.64% of the CDA variance in global fibrosis in younger mice treated with doxorubicin or combined therapy, respectively (**Fig. 6B**.)

**Figure 6.**
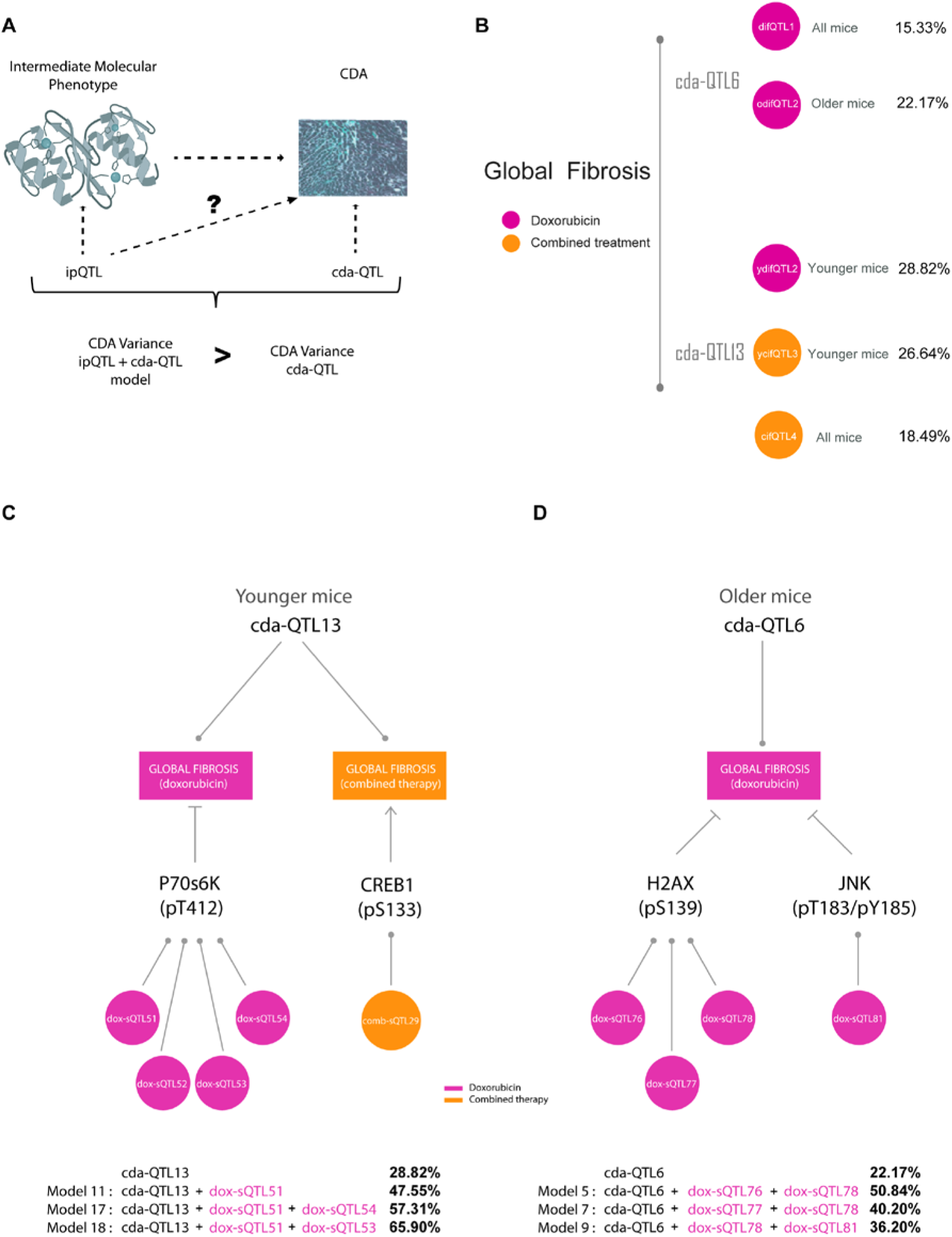
Percentage of global fibrosis explained by cda-QTL6 or cda-QTL13 and genetic models. **A**) Genetic models between ipQTLs and cdaQTL are generated to assess whether an ipQTL contributes to the phenotypic variation of CDA. If the variability of the model significantly exceeds that of the cdaQTL, the ipQTL would contribute to the phenotypic variation of CDA and susceptibility. **B**) Above, the diagram shows the percentage of global fibrosis explained by cda-QTL6 after treatment with doxorubicin (red). The circles on the right show the name of the cda-QTL6 under these conditions, difQTL1 (doxorubicin-induces fibrosis QTL1), which appears linked to global fibrosis in all mice, and odifQTL2 (old mice difQTL2), which seems to be associated with global fibrosis in all mice. Below is the percentage of global fibrosis explained by cda-QTL13 following treatment with doxorubicin (red) or combined therapy (yellow). The circles on the right show the other names of the cda-QTL13 for those conditions: ydifQTL2 (young mice doxorubicin-induced fibrosis QTL2), ycifQTL3 (young mice combined therapy-induced fibrosis QTL3), and cifQTL4 (combined therapy-induced fibrosis QTL3), the latter linked to global fibrosis in all mice. **C**, **D**) we chose cda-QTL6 and cda-QTL13 to assess whether ipQTLs combined with these cda-QTLs could account for more of the CDA trait variation than that justified exclusively by cda-QTL6 and cda-QTL13. Thus, we selected cda-QTL6 and cda-QTL13 as examples to generate the genetic models. Schemes illustrating the components used to develop the genetic models (Table 1). **C**) In this case, the objective was to evaluate whether genetic models can explain more global cardiac fibrosis variance after doxorubicin treatment in young mice than cda-QTL13. To this end, the two intermediate molecular phenotypes associated with global fibrosis in young mice after treatment with doxorubicin were P70S6K(pT412), and CREB1(pS133) (Table 1A and **Supplemental_Table_S3.xls**) and the ipQTLs linked with them are shown in Table 1A and **Supplemental_Table_S5.xls**. The genetic models were developed with cda-QTL13 and four combinations of the ipQTLs linked to P70S6K(pT412) (dox-ipQTL51, dox-ipQTL52, dox-ipQTL53, and dox-ipQTL54), and one combination, comb-ipQTL29, linked to CREB1(pS133). The phenotypic variability explained by cda-QTL13 alone and the significant increase in those genetic models in which it improved are indicated below the diagram (Table 1B). **D**) Scheme to illustrate the same results as panel C, but for the case of cda-QTL6 and global fibrosis in old mice. Panels C and D correspond to Table 1.

We then assessed whether ipQTLs linked to myocardium molecules increased the amount of global heart fibrosis explained by cdaQTL6 in old mice and cdaQTL13 in young mice. The intermediate molecular phenotypes that were correlated with global fibrosis under these conditions and their ipQTLs are shown in **Fig. 6C, D**, and **Table 1A**. We examined all the viable genetic models with cdaQTL6 or cdaQTL13 and the ipQTLs linked to the intermediate molecular phenotypes associated with global fibrosis(Broman et al. 2003) (**Supplemental_Table_S8.xls**).

**Table 1.** Genetic models. **A)** QTLs used to develop the genetic models, **B)** Improvement of the CDA variation explained by cdaQTL6 or cda-QTL13 with ipQTLs integrated into genetic models. Genetic models combining ipQTLs linked to intermediate molecular phenotypes increase the proportion of phenotypic variation of global fibrosis explained by cdaQTL13 in younger mice and cda-QTL6 in older mice treated with doxorubicin. Only models that improved the fibrosis phenotypic variability explained are included (**Supplemental_Table_S8.xls**). Genetic models were generated by the Fitqtl function (r/qtl package). This table is related to Fig. 6C, D.

| A) QTLs and CDA conditions used in the genetic models |  |  |  |  |  |
| --- | --- | --- | --- | --- | --- |
| cda-QTL | Therapy type | Mouse age group | Pathophenotype of CDA | (a) Molecular Intermediate Phenotypes Associated with Global Fibrosis | (b) ipQTLs |
| cda-QTL6 | Doxorubicin | Old Mice | Global Fibrosis | γH2AX(S139) | dox-ipQTL76 |
|  |  |  |  |  | dox-ipQTL77 |
|  |  |  |  |  | dox-ipQTL78 |
|  |  |  |  | JNK(T183/Y185) | dox-ipQTL81 |
|  |  |  |  | miR200b-3p | N.I. |
| cda-QTL13 | Doxorubicin | Young Mice | Global Fibrosis | p70S6K(T412) | dox-ipQTL51 |
|  |  |  |  |  | dox-ipQTL52 |
|  |  |  |  |  | dox-ipQTL53 |
|  |  |  |  |  | dox-ipQTL54 |
|  | Combined Therapy | Young Mice | Global Fibrosis | CREB(S133) | comb-ipQTL29 |

| B) Improvement of the global fibrosis variation explained by cda-QTL6 and cda-QTL13 with ipQTLs in genetics models |  |  |  |  |  |
| --- | --- | --- | --- | --- | --- |
| Basal effect / Model effect |  | Model components | LOD score | Fibrosis variation (%) | P - value |
| cda-QTL13 | Basal effect | n.a. | 2.38 | 28.82 | n.a. |
| Models with cda-QTL13 in young mice | Model 11 | cda-QTL13 * dox-ipQTL51 | 3.78 | 47.55 | 0.0006 |
|  | Model 17 | cda-QTL13 * dox-ipQTL51 * dox-ipQTL54 | 4.99 | 57.31 | 0.002 |
|  | Model 18 | cda-QTL13 * dox-ipQTL51 * dox-ipQTL53 | 6.3 | 65.90 | 0.0001 |
| cda-QTL6 | Basal effect | n.a. | 2.29 | 22.17 | n.a. |
| Models with cda-QTL6 in old mice | Model 5 | cda-QTL6 * dox-ipQTL76 * dox-ipQTL78 | 6.63 | 50.84 | 0.00007 |
|  | Model 7 | cda-QTL6 * dox-ipQTL77 * dox-ipQTL78 | 4.8 | 40.20 | 0.0024 |
|  | Model 9 | cda-QTL6 * dox-ipQTL78 * dox-ipQTL81 | 4.2 | 36.20 | 0.0075 |

With respect to global heart fibrosis in young mice, we observed that the phenotypic variation due to cda-QTL13 increased after including some of the ipQTLs associated with P70S6K in genetic models (**Fig. 6C** and **Table 1B**). The ipQTLs linked to P70S6K levels in young mice were located on Chr. 3 (dox-ipQTL51), Chr. 4 (dox-ipQTL52), Chr. 9 (dox-ipQTL53), and Chr. 17 (dox-ipQTL54) (**Table 1A**). The variance in global fibrosis explained by cda-QTL13 was 28.82%. This increased to 47.55% when considering the dox-ipQTL51 (model 11), to 57.31% when including the dox-ipQTL51 and dox-ipQTL54 (model 17), and to 65.9% for dox-ipQTL51 and dox-ipQTL53 (model 18) (**Fig. 6C** and **Table 1B**).

As indicated, the criteria for choosing intermediate molecular phenotypes of CDA were based on the evidence from previous studies. In the case of P70S6K, protective and anti-protective effects after treatment with doxorubicin have both been described(Xu et al. 2012; Yu et al. 2013; Lee et al. 2015). We confirmed its role in CDA through functional *in vitro* studies. Human-induced pluripotent stem cell-derived cardiomyocytes (hi-PSC-CMs) are used to ensure the involvement of genes in cardiotoxicity at a functional level(Sharma et al. 2018). Hence, as an example, we demonstrated that downregulating *RPS6KB1* levels with siRNA in hiPSC-CMs increases their sensitivity to doxorubicin, confirming the role of P70S6K as an intermediate molecular phenotype of CDA (**Supplemental_Fig**_**S3.pdf.**)

Similarly, concerning global heart fibrosis in old mice, cdaQTL6 explained 22.17% of the phenotypic variation. This value was higher when ipQTLs associated with myocardium levels of ɤH2AX and pJNK were included in the models (**Fig. 6D** and **Table 1**). The amount of phenotypic variation in CDA explained by genetic determinants directly linked to CDA (cdaQTLs) was boosted using ipQTLs linked to intermediate molecular phenotypes associated with CDA, implying that the genetic determinants controlling the intramyocardial levels of these intermediate molecular phenotypes contribute to the phenotypic variance of CDA. However, as they are QTLs directly linked to the CDA, we can deduce that they are the source of some of the missing CDA heritability. This large amount of phenotypic variation elucidated can be explained by the simplicity of the backcross model, with a more limited genetic diversity than human populations (Buchner and Nadeau 2015).

### Genes encoding intermediate molecular phenotypes associated with myocardium damage in mice can be genetic determinants of CDA in patients

The previous analyses showed that ipQTLs linked to the levels of intermediate molecular phenotypes of CDA increase the proportion of phenotypic variation explained by cda-QTLs. However, the levels of these intermediate phenotypes associated with CDA also depend on the regulatory regions of the genes encoding the intermediate phenotypes themselves. Given this, we can presume that all of them, the regulatory regions in *cis* with the gene encoding the intermediate phenotype, and the ipQTLs, mostly in *trans*, may contribute through the levels of the intermediate molecular phenotype to CDA variation. We have not identified the ipQTL driver, but it is reasonable to propose that the genes known to encode the intermediate molecular phenotypes of CDA probably can be genetic determinants of CDA susceptibility (**Fig. 1B, C**).

We evaluated the extent to which allelic forms of the genes encoding intermediate molecular phenotypes associated with myocardium damage in mice could be genetic determinants of CDA in three cohorts of cancer patients treated with anthracyclines. CDA was evaluated by echocardiography in a cohort of women with breast cancer and another with pediatric cancer; in the third cohort, the CDA was evaluated by cardiac magnetic resonance (CMR), considering an LVEF reduction of 5% or more during the first six months or throughout the complete follow-up. We only evaluated SNPs from those genes that encoded molecules associated with CDA in mice, testing the most probably genetic model (**Fig. 3** and **Supplemental_Table_S3.xls**). Several single nucleotide variants (SNVs) were associated with susceptibility to CDA in patients, some of which were noted in more than one cohort (**Supplemental_Table_S9.xls**).

Subsequently, we used the two largest cohorts, formed by breast cancer patients and pediatric patients, to generate two polygenic risk scores. Thus, each cohort was divided into a training set (80%) and a testing set (20%). In the training set, after bootstrapping 100 times, a series of SNPs associated with susceptibility to CDA were identified. Subsequently, two different risk models were obtained using the restrictive LASSO regression system (**Fig. 7**). Interestingly, the SNPs that were part of the risk scores belonged to the same five genes in both models. Indeed, the CDA model in adults consisted of SNPs of *AKT1, STAT3, TP53, MAPK8, MAPK11*, and *RelA/P65*; the pediatric model consisted of SNPs of the same first five genes.

**Figure 7.**
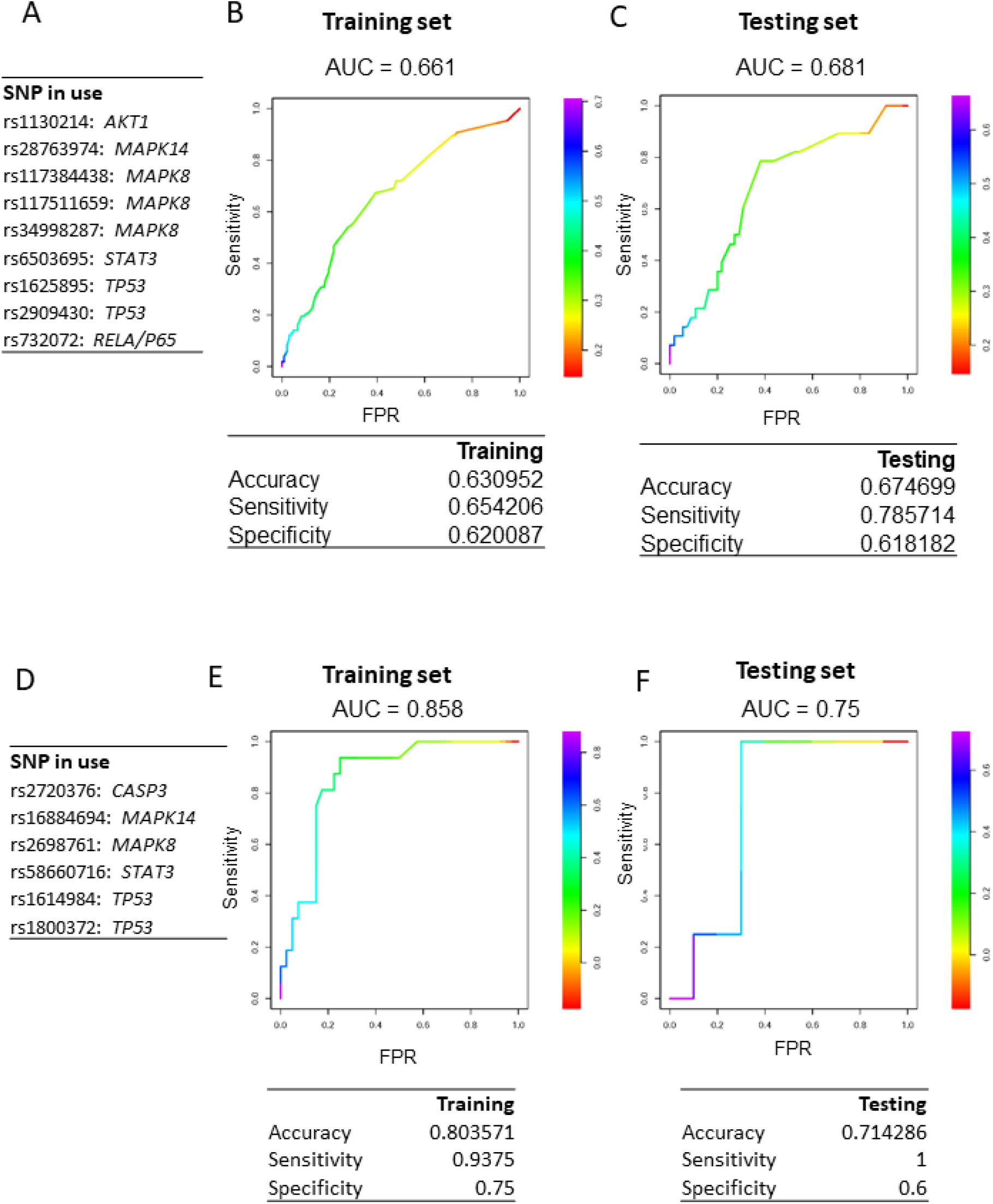
Genetic models of CDA risk generated by bootstrapping 100 times and LASSO multivariate regression. **A-C**) CDA risk model for the cohort of breast cancer patients. The genetic model (A) and the ROC curves of the training set (B) and the testing set (C) are shown. **D-E**) Genetic risk model for the pediatric patient cohort. The genetic model (D) and the ROC curves for the training set (E) and the testing set (F) are shown. Cohorts were split at 80% for the training and 20% for the testing set. FPR, false positive rate.

Our results highlight the value of genetic mouse models as tools for identifying the intermediate phenotypes that contribute to the variation of CDA and the use of the genes encoding them as potential susceptibility markers.

## Discussion

Chronic CDA is a common side effect that can be very severe and affect the continuity of chemotherapy treatment (Caron and Nohria 2018). CDA susceptibility varies considerably among patients (Grenier and Lipshultz 1998), but significant efforts have been made, through the use of genetic markers, to identify the patients who are susceptible to developing chronic CDA(Duan et al. 2007). Predisposition to chronic CDA has a strong genetic component and, as a complex disease or trait, it is, by definition, polygenically inherited (Leong et al. 2017). The genetic elements of complex characters are difficult to identify; there are substantial discrepancies between the proportion of phenotypic variance expected to arise from genetic causes (expected heritability) and the heritability explained by identified DNA sequencing variants (DSVs). The difference between them is known as missing heritability (Manolio et al. 2009). CDA, being a complex trait, has an unknown degree of missing heritability, making it challenging to identify most of its genetic components.

The use of intermediate phenotypes to identify a part of the genetic component of complex traits has been proposed in psychiatric disorders (Gottesman and Gould 2003) and extended to other fields(Blanco-Gomez et al. 2016). It has been suggested that identifying the genetic determinants associated with intermediate phenotypes essential for complex-trait pathogenesis could help identify a part of the missing heritability and yield genetic markers for predicting complex disease susceptibility (Blanco-Gomez et al. 2016). Part of the genetic component of the missing heritability could be explained by the genetic determinants, such as QTLs, that are linked to intermediate phenotypes involved in the pathogenesis of complex traits. Indeed, a highly influential QTL can be simultaneously detected at the intermediate phenotype and primary complex trait levels, reflecting a process known as mediated pleiotropy (Solovieff et al. 2013). It has been suggested that the QTLs that cannot be detected as mediated pleiotropy are part of the missing heritability because they are too weak to be revealed by genetic analysis at the complex-trait level (Castellanos-Martin et al. 2015; Blanco-Gomez et al. 2016).

The network of intermediate phenotypes at the systemic, tissue, cellular and molecular levels that determines the pathogenesis of a complex trait is regulated by a multitude of genes acting at all those levels. This scenario coincides with the omnigenic model, involving a series of gene networks with core genes and many peripheral genes, that has recently been proposed to explain missing heritability(Boyle et al. 2017; Liu et al. 2019). It is inferred from this model that the complete heritability of a complex trait is controlled by most of the genome (Loh et al. 2015; Khera et al. 2018; Liu et al. 2019). The source of most heritability is genes with so little effect that many are difficult, or even impossible, to identify, no matter how many individuals are studied.

From a practical point of view, the challenge is to find a way to determine the genetic markers of susceptibility to a complex trait. The focus on the genetic determinants associated with the variation of intermediate phenotypes (which, in turn, are associated with a complex trait) makes it possible to identify essential genes that may be involved in the missing heritability and that can be susceptibility markers of diseases of complex genesis. In this sense, the approach proposed in this study could be adopted as a general strategy for better identifying genetic markers of high-prevalence complex diseases, e.g., type II diabetes, autoimmune diseases, thrombosis, cardiovascular diseases, sporadic cancers, and CDA (Maher 2008).

The initial search for genetic determinants associated with intermediate phenotypes can be simplified in models of limited genetic variability, such as crosses of genetically homogeneous mouse strains (Hunter and Crawford 2008; Quigley and Balmain 2009). It is difficult to determine the polygenic component of complex human population traits because of their genetic complexity and sophisticated interaction with the environment(Hunter and Crawford 2008). Identifying genes with a weak effect in human studies using techniques such as GWAS is complicated because enormous sample sizes are required to demonstrate statistical significance. In addition, the massive amount of multiple testing supposed by the analysis of millions of SNPs dramatically reduces statistical power, especially when trying to locate variants of common genetic variants of weak effect. However, quantifying the phenotypic variation of complex traits, under controlled environmental conditions, in a simplified genetic model consisting of crosses between genetically homogeneous strains of mice can guide the choice of candidate genes and pathways to be tested in human populations. Identifying candidate genes in this simplified genetic model reduces the number of genetic variants that need to be considered(Quigley and Balmain 2009). We think this strategy makes it possible to identify genetic markers of complex traits without carrying out studies in thousands of patients because evaluating intermediate molecular phenotypes in the simplified genetic model in mice enables the selection of candidate genes. One of the main limitations of GWAS is the difficulty of subsequent validation in other populations(Wray et al. 2013). Thus, to confirm in humans the candidate genes identified in the genetically heterogeneous cohort of mice, we think it could be more effective to use several cohorts of cancer patients with different conditions than a larger, though more homogeneous cohort, as we have done here with several distinct patient cohorts treated with anthracyclines. We think finding the association of genetic markers with CDA simultaneously in several of these other conditions increases the possibility of them being genuine and of subsequent utility.

This work has identified part of the genetic component linked to intermediate molecular phenotypes associated with CDA in a mouse backcross model, which helped identify some genetic elements related to the CDA susceptibility itself. Since anthracyclines exert their toxicity by damaging DNA, CDA intermediate phenotypes would differ in terms of the molecular pathways involved in DNA damage response or in the signaling pathways, such as AKT (Ichihara et al. 2007) and P38MAPK (Kang et al. 2000), that promote or protect from heart damage by anthracyclines (Ghigo et al. 2016). We also hypothesize that other CDA intermediate phenotypes can be molecules involved in heart diseases and cardiotoxicity, such as cell signaling pathways (Ghigo et al. 2016), miRNAs (Ruggeri et al. 2018), and telomere length (De Angelis et al. 2010). Multiple regression models allowed us to pinpoint which of these intermediate molecular phenotypes, determined in mouse myocardium, are best able to explain the phenotypic variation in the CDA. We then evaluated the extent to which the ipQTLs associated with these intermediate molecular phenotypes account for chronic CDA.

Although ipQTLs were not directly linked to CDA, we used genetic models to demonstrate that ipQTLs allow more of the phenotypic variability of QTLs directly linked to CDA to be explained, thereby showing that these ipQTLs account for part of the missing heritability of the CDA. They help control in *trans* the levels of intermediate protein phenotypes located in the myocardium. One of the variables that most strongly influences the activity of a molecule is its level, and this depends on factors in *cis*, the most important of these being the gene sequences encoding the molecule and various elements in *trans*. However, we cannot rule out the possibility that any of the ipQTLs identified may act at other levels, for example, by contributing to the control of the proteins’ phosphorylation. We have used ipQTLs in *trans* to demonstrate that genetic determinants linked to the levels of the molecule contribute to the phenotypic variability of the complex trait. Although we do not know which genes drive the effects of ipQTLs, we know the gene encoding the intermediate protein phenotype in *cis*. So, if genetic variants of these genes do determine the levels of the protein intermediate phenotype, they will also contribute to the heritability of the complex trait. For this reason, we looked for variants of these genes to evaluate in the human population.

An enormous number of intermediate phenotypes affect the pathogenesis and variation in a complex trait like CDA, so it is unsurprising that many of the contributing genetic determinants cannot be detected among those of the central phenotype, for which reason they are responsible for much of the missing heritability (Blanco-Gomez et al. 2016). Furthermore, the myriad interactions between intermediate phenotypes and the abundance of QTLs associated with them make it unlikely that the sources of missing heritability of a complex phenotype, including CDA, could ever be accounted for completely. Indeed, these intermediate phenotype interactions at different levels may involve most of the genome (Liu et al. 2019).

### Conclusions

A genetically heterogeneous cohort of mice was used to identify the genetic component of proteins whose levels in the myocardium are associated with histopathological damage after chemotherapy, and thereby to reveal some of the missing genetic elements linked to CDA in mice and humans. Identifying genetic and molecular factors responsible for the increased risk of CDA will eventually improve our ability to predict and prevent CDA. Our results suggest that, in general, the proposed strategy facilitates the identification of susceptibility markers of complex diseases. The genetic markers identified could also help identify patients at high risk of developing CDA, enabling personalized patient management and optimized individualized chemotherapy to reduce the risk of severe adverse drug reactions.

## Methods

### Patients

The association of genetic variants with CDA was evaluated in four patient cohorts previously published by some of us. In the first three cohorts, comprising 71 anthracycline-treated pediatric cancer patients (Ruiz-Pinto et al. 2017) (Paediatric Cohort) and 420 breast cancer patients (Breast Cancer Cohort) (Vulsteke et al. 2015), cardiac function was assessed by echocardiography to evaluate the left ventricular ejection fraction (LVEF) or left ventricular fractional shortening (LVFS). In the third cohort, cardiac magnetic resonance (CMR) was carried out in 24 cancer patients (CMR cohort) (Barreiro-Perez et al. 2018) at baseline and after every two cycles of a regular course of anthracycline therapy. All patients received anthracyclines in their treatment. Their clinical features have already been published (Vulsteke et al. 2015; Ruiz-Pinto et al. 2017; Barreiro-Perez et al. 2018; Ruiz-Pinto et al. 2018). Following the Declaration of Helsinki, we obtained the Bioethics Committee’s permission and the informed consent of the patients or their relatives in the case of pediatric patients. The CMR study is described below and was approved by the University Hospital of Salamanca’s Institutional Ethics Review Board.

### Cardiac magnetic resonance: acquisition and analysis

Cardiac magnetic resonance (CMR) examinations were conducted with a Philips 1.5-Testa Achieva whole-body scanner (Philips Healthcare) equipped with a 16-element phased-array cardiac coil and fully installed and managed by the Cardiology Department at the University Hospital of Salamanca (Barreiro-Perez et al. 2018). The imaging protocol always included a standard segmented cine steady-state free-precession (SSFP) sequence to provide high-quality anatomical references. The imaging parameters for the SSFP sequence were: 280 × 280 mm field of view, 8 mm slice thickness with no gap, 3 ms repetition time, 1.50 ms echo time, 60° flip angle, 30 cardiac phases, 1.7 × 1.7 mm voxel size and a single excitation. CMR images were analyzed using dedicated software (MR Extended Work Space 2.6, Philips Healthcare, Netherlands) by two observers experienced in CMR analysis and blinded concerning time-point allocation and patient identification.

### Mouse generation and chemotherapy

We generated a genetically heterogeneous mouse cohort by backcrossing two inbred strains. We crossed a breast cancer-resistant mouse strain, C57BL/6 (hereafter C57), with a susceptible strain, FVB/J, to generate F1 mice. Later, the non-transgenic F1 mice were crossed with *FVB/N-Tg(MMTVneu)202Mul/J* transgenic mice (hereafter FVB), carrying the *Avian erythroblastosis oncogene B2/Neuroblastoma-derived* (*ErbB2/cNeu*) protooncogene, expressed under the mouse mammary tumor virus (MMTV) promoter (MMTV-*Erbb2/Neu* transgene) and allowed to develop breast cancer(Guy et al. 1992).

Each mouse from the backcross cohort carried a unique combination of alleles from the two strains (FVB and C57) in variable proportions. In this combination, the genetic component from the FVB strain was predominant since it was the one used to generate the backcross with the F1 mice. FVB alleles can be homozygous or heterozygous, while the C57 component is reduced and heterozygous when present. The cross was designed to enrich the alleles for susceptibility to breast cancer in the cohort. The mice were administered chemotherapy once they had developed breast cancer under isofluorane anesthesia. Mice were euthanized by CO2 when the tumors were bigger than 15 mm or showed signs of suffering. We evaluated cardiotoxicity in 164 mice: 130 F1BX, 18 FVB, and 16 F1 (the latter having been generated after crossing FVB transgenic mice with C57). FVB transgenic mice were obtained from the Jackson Laboratories, and wild-type FVB/N and C57BL/6 mice were purchased from Charles River.

All mice were housed in ventilated filter cages in the Animal Research Facility of the University of Salamanca under specific-pathogen-free (SPF) conditions and fed and watered *ad libitum*. One group (N = 87) was treated with doxorubicin every 10 days with a dose of 5 mg/kg, and another group (N = 77) received the combined therapy of doxorubicin (Pfizer) (5 mg/kg) plus docetaxel (Sanofi Aventis) (25 mg/kg), administered intraperitoneally every 10 days. The drug doses used mimicked the clinically relevant drug concentration (Rottenberg et al. 2007). Mice received four therapy cycles or five if the chemotherapy was well-tolerated. Once the treatment had finished, the mice’s evolution and tumor development were assessed for 2 months. Necropsies were then performed, and the heart and other tissues were collected. All practices were previously approved by the Institutional Animal Care and Bioethics Committee of the University of Salamanca and conformed to the guidelines from Directive 2010/63/EU of the European Parliament on animals’ protection for scientific purposes.

### Mouse genotyping

Briefly, DNA was extracted from the tail by the phenol-chloroform method. DNA concentrations were measured with a Nanodrop ND-1000 Spectrophotometer, and the PicoGreen double-stranded quantification method (Molecular Probes, Thermo Fisher Scientific Inc., Waltham, MA USA) was used for genotyping. Genome-wide scanning was carried out at the Spanish National Centre of Genotyping (CeGEN) at the Spanish National Cancer Research Centre (CNIO, Madrid, Spain). The Illumina Mouse Medium Density Linkage Panel Assay was used to genotype 130 F1BX mice at 1449 single nucleotide polymorphisms (SNPs). Genotypes were classified as FVB/FVB (F/F) or FVB/C57BL/6 (F/B). Ultimately, 806 SNPs were informative from the FVB and C57BL/6 mice; the average genomic distance between these SNPs was 9.9 Mb. The genotype proportion among the F1BX mice was normally distributed.

### Heart-tissue processing and CDA quantification

Hearts were fixed in 4% paraformaldehyde (Scharlau FO) for 24 hours and then processed in an automatic system (Shandon Excelsior, Thermo). The subsequent samples were sectioned, embedded in paraffin, and stained with hematoxylin-eosin with a standard protocol or the Masson Trichrome Goldner kit (Bio-Optics) to evaluate the cardiac fibrosis cardiomyocyte area. We automatically quantified heart fibrosis and the average area of myocardial fibers as pathophenotypes of CDA using the Ariol slide scanner to avoid intra- and inter-observer deviations. Histopathological damage was measured in the subendocardium and subepicardium from five randomly chosen regions of each sample.

### Protein extraction

Approximately 10-15 mg of frozen cardiac tissue were homogenized using the FastPrep Homogenizer system (FP120, Bio 101 Thermo Savant) and ceramic beads (Precellys Lysing Kit CkMix, Precellys) in lysis buffer (Lysis Buffer 1X, Milliplex) to which a cocktail of protease inhibitors (Roche Complete Mini) and phosphatase inhibitors (PhosSTOP EASYpack, Roche) was added. The quantification of signaling proteins and other intermediate molecular phenotypes is described in the supplementary methods.

### hiPSC-CMs infection and viability analysis

The supplementary methods describe the generation of human-induced pluripotent stem cell-derived cardiomyocytes (hi-PSC-CMs) and lentiviral infection.

### Mouse QTL genetic analyses and genetic models

Linkage analysis was carried out using interval mapping with the expectation-maximization (EM) algorithm and R/QTL software. The criteria for significant (lod score > 3) and suggestive (lod score > 1.5) linkages for single markers were chosen based on the findings of Lander and Kruglyak(Lander and Kruglyak 1995). In the QTL results tables, the cXX.loc.XX markers do not refer to real SNPs but instead to genetic locations where the conditional genotype probabilities for the EM algorithm were calculated using the *calc.genoprob* function in R/qtl, with a step of 2.5 and an error.prob of 0.001. Various QTL models were developed in which the *fitQTL* function was used with Haley-Knott regression in R/qtl to fit and compare their LOD scores and the percentage of the explained variance (Broman et al. 2003). QTLs were chosen for inclusion in the final model only if they demonstrated a significant additive or interaction effect (*p* < 0.05), determined by a “drop-one-QTL-at-a-time” analysis, which evaluates the impact of single QTLs or interactions.

### Human genetic analysis

The existence of associations of CDA, measured by echocardiography or CMR, with SNVs was evaluated in the four patient cohorts. We looked for alleles encoding proteins whose levels in the myocardium were associated with CDA in mice. SNVs were detected on the Infinium™ Global Screening Array-24 v2.0 BeadChip. Data were imputed using the Michigan Imputation Server with Minimac4(Das et al. 2016). After retrieving the data, all markers with R^2^ < 0.7 were removed from the analysis before proceeding further. Data were analyzed in R v3.6.0(1). The SNPassoc package v2 was used to explore associations between mutations. Employing the “association” function, we performed case/control analysis (Chi-square test) of all the possible genetic models (codominant, dominant, recessive, overdominant, and log-additive) to examine the associations between phenotypes and input mutations.

### Prediction of CDA in Human Cohorts

Each human cohort was first randomly partitioned into 80% training set and 20% testing set. During model construction, logistic regression was deployed in a bootstrapping strategy with a fixed sampling rate (80% evaluation and 20% validation) and many iterations (100). After bootstrapping, the SNPs, significantly associated with CDA in both evaluation and validation data for at least 5 times across 100 iterations, were selected to construct the Least Absolute Shrinkage and Selection Operator (LASSO) regression model on the 80% training set. After training, the best cutoff on the receiver operating characteristic (ROC) curve was optimized based on the maximum Youden’s index (sensitivity+specificity-1), and the LASSO models, as well as the corresponding optimal cutoff, were applied on the 20% testing set for independent evaluation of the model performance.

For addicinal information, see also **Supplemental_Methods.pdf**.

## Data access

This published article and its supplemental information files include most of the data generated and analyzed in this study. Related metadata underlying the findings are available as additional datasets in the public repository DIGITAL.CSIC http://hdl.handle.net/10261/239215. The other human genetic and clinical data are available upon reasonable request from those of us who are the corresponding authors of previously published manuscripts.

<u>Supplementary Datasets that are available in the DIGITAL_CSIC repository are:</u>

1. Folder-1: datasets related to the quantification of CDA pathophenotypes.
2. Folder-2: datasets associated with the quantification of intermediate molecular phenotypes.
3. Folder-3: datasets related to mouse genotyping.
4. Folder-4: datasets associated with LOD scores of ipQTLs.
5. Folder-5: datasets associated with LOD scores of cdaQTLs.
6. Folder-6: datasets related to patients.

## Acknowledgments

JPL’s lab is sponsored by Grant PID2020-118527RB-I00 funded by MCIN/AEI/10.13039/501100011039; Grant PDC2021-121735-I00 funded by MCIN/AEI/10.13039/501100011039 and by the “European Union Next Generation EU/PRTR,” the Regional Government of Castile and León (CSI144P20), the Carlos III Health Institute (PIE14/00066). AGN laboratory and human patients’ studies are supported by an ISCIII project grant (PI18/01242). The Human Genotyping unit is a member of CeGen, PRB3, and is supported by grant PT17/0019, of the PE I+D+i 2013-2016, funded by ISCIII and ERDF. SCLl was the recipient of a Ramón y Cajal research contract from the Spanish Ministry of Economy and Competitiveness, and the work was supported by MINECO/FEDER research grants (RTI2018-094130-B-100). CH, was supported by the Department of Defense (DoD) BCRP, No. BC190820; and the National Cancer Institute (NCI) at the National Institutes of Health (NIH), No. R01CA184476. Lawrence Berkeley National Laboratory (LBNL) is a multi-program national laboratory operated by the University of California for the DOE under contract DE AC02-05CH11231. The Proteomics Unit belongs to ProteoRed, PRB3-ISCIII, supported by grant PT17/0019/0023 of the PE I + D + I 2017-2020, funded by ISCIII and FEDER. RCC is funded by fellowships from the Spanish Regional Government of Castile and León. NGS is a recipient of an FPU fellowship (MINECO/FEDER). hiPSC-CM studies were funded in part by the “la Caixa” Banking Foundation under the project code HR18-00304” and a Severo Ochoa CNIC Intramural Project (Exp. 12-2016 IGP) to JJ. We thank Isabel Ramos and Marina Jiménez for their help in the Animal House Facility, Elena Alonso for technical assistance, and Yolanda Gómez-Vecino for helping to design some of the figures. We thank Emma Keck and Phil Mason for the English language support.

